# Reintroductions backfire by destabilising food webs and triggering further extinction cascades

**DOI:** 10.1101/2025.08.28.672799

**Authors:** Thomas F. Johnson, Alain Danet, Toby Haefling, Andrew P. Beckerman

## Abstract

Conservation interventions like reintroduction are considered vital to bending the curve of biodiversity loss^1^, with the potential for multiplicative benefits that not only reduce extinction pressure on the reintroduced species, but also restore wider community dynamics and ecosystem functions^2,3^. Its growing popularity reflects these perceived benefits^4^. Reintroduction success is usually judged only by the survival of the released species^4^, but there is no guarantee that the wider community will recover. Using simulated food webs, we show that reintroductions can frequently have unintended negative consequences: triggering extinction cascades, reducing biomass, and destabilising communities. These perverse impacts are unlikely to be detected as reintroduction success is often measured solely by the ability for reintroduced species to survive - which our models suggest is likely - introducing a risk that conservation action superficially appears successful but is actually further depleting our ecosystems.

---

The Kunming-Montreal Global Biodiversity Framework aims to protect and restore ecosystems, and the vital services they provide^5^. This is an ambitious and urgent goal, set against a backdrop of intense anthropogenic pressures from land use, climate change, species invasions, overexploitation, and pollution^6^. Reducing these threats is essential to halting biodiversity loss, but alone it is unlikely to guarantee recovery. To bend the curve of biodiversity loss towards recovery, degraded landscapes and ecological communities often need to undergo active restoration and conservation^1^, helping to reverse declines and stabilise population and community dynamics.

Interventions such as species reintroductions are an increasingly important tool for rebuilding degraded ecological communities and are being adopted more widely as a conservation strategy ^4^. Traditionally, reintroduction success has been judged by whether the reintroduced species can survive and reproduce^4^, which hinges on habitat and community conditions remaining sufficiently intact to support positive population growth. Recent influential studies, however, argue that reintroductions can achieve more than safeguarding the focal species - they may also restore broader community dynamics and critical ecosystem functions^2,7^. Yet these wider benefits are rarely quantified^4^, and it remains uncertain how consistently they are realised.

Although it is often assumed that reintroducing a species will automatically benefit its community, this is far from guaranteed. First, the very pressures that caused the focal species’ extirpation may also have driven widespread losses in other species, degrading the community as a whole. Second, the disappearance of the focal species itself can trigger extinction cascades, where the loss of a single species disrupts trophic interactions or shifts competitive hierarchies^8^, and can redistribute interaction strengths among those species that remain^9^. In such contexts, the community has changed to such a degree that the reintroduced species may fail to establish, and worse, its return could further destabilise the community through processes such as competitive exclusion. Understanding when reintroductions are likely to succeed - and recognising that past extirpations can reshape communities in multiple ways - is therefore crucial if reintroduction is to be a reliable tool for bending the curve of biodiversity loss.

Acknowledging both the potential benefits and risks of reintroductions is critical to ensuring success across ecological scales (from population to community and ecosystem). However, empirical studies assessing community-level consequences remain rare^4^, as they require long-term monitoring of multiple species. In the absence of such evidence, theoretical modelling offers a powerful tool to test reintroduction scenarios and evaluate both benefits and risks. Here, we use a multi-species bioenergetic food-web model^10,11^ to explore how predator extirpation and reintroduction reshape ecological communities. We focus on top predators, which are often considered keystone species that regulate community dynamics^12,13^, but which are also among the most frequently extirpated^14,15^ and often targets for reintroduction.

We first investigate how community richness, biomass, evenness and temporal stability change after predator loss. We then reintroduce the predator into the community, examining how community intactness (i.e. the proportion of the intact community that remains) influences the probability of re-establishment. Finally, we assess whether reintroduction produces unintended consequences for the community, such as triggering further extinctions, redistributing biomass, or altering stability and evenness. We explore these outcomes across a standard range of food-web structural characteristics - including species richness, connectance (link density/complexity), and consumer–resource body-size ratios. Ultimately, our aim is to identify the conditions under which reintroductions succeed in restoring communities to more intact states, versus those where they risk imposing further destabilisation.

## Predator extirpation triggers extinction cascades, biomass change and loss of stability

The loss of a predator can have substantial impacts on community composition. Predator removal consistently led to secondary extinctions, regardless of food-web structure, complexity, or predator traits such as size and diet breadth (Figure 1A). It also reshaped biomass distribution: when the lost predator was relatively small, occupied a high trophic position, and had a narrow diet breadth, community biomass increased substantially (Figure 1B), while stability remained largely unchanged (Figure 1C). In contrast, the removal of a large predator with a lower trophic position and broad diet breadth produced only marginal biomass gains but caused a marked decline in stability. In nearly all cases, predator loss reduced community evenness, with trophic cascades shifting biomass toward lower trophic levels (Figure 1D).

**Figure 1.**
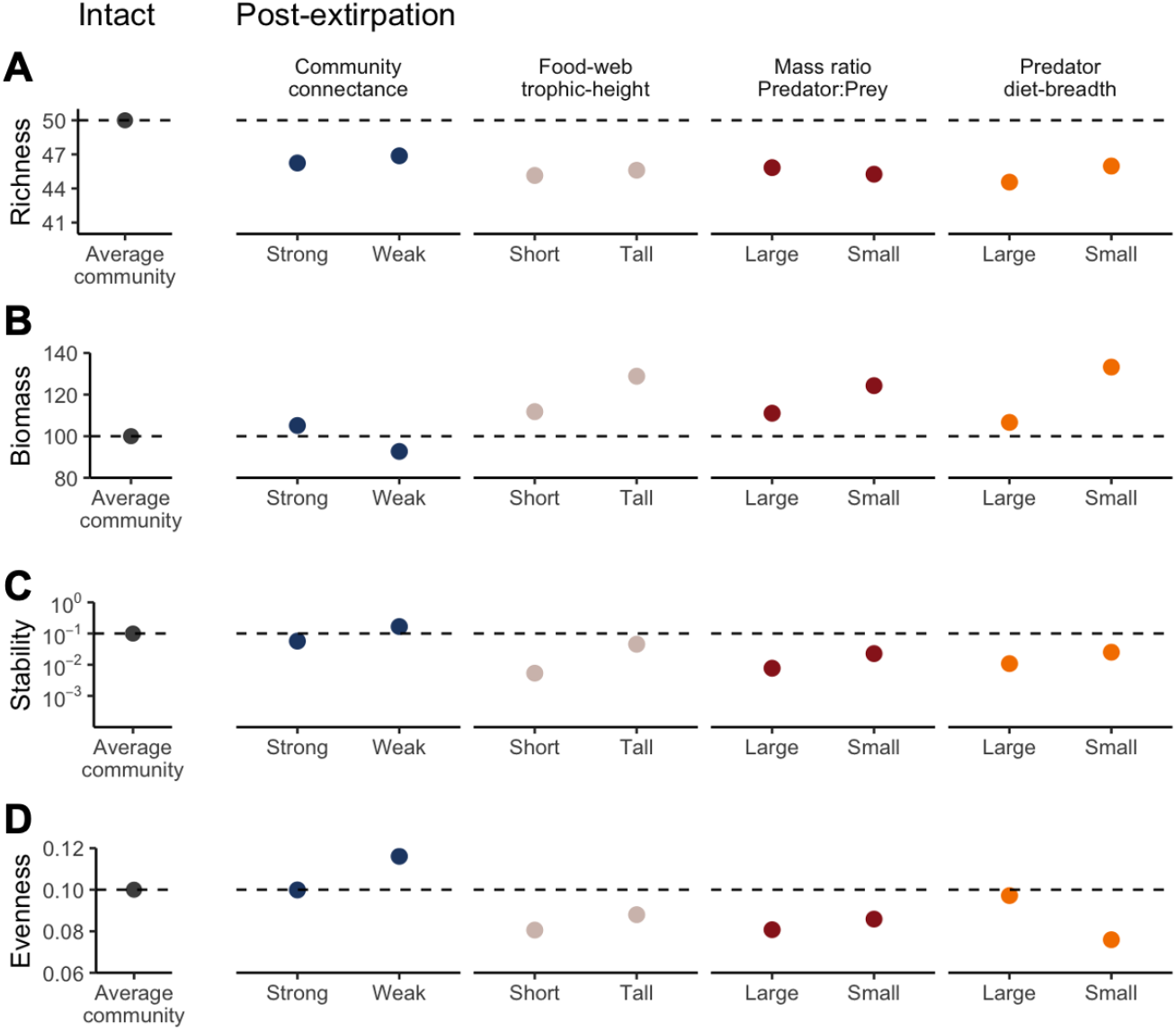
Community responses to the extirpation of the top predator. Change in community richness (A), biomass (B) stability (C) and evenness (D) in a hypothetical ‘average’ community under four post-extirpation scenarios each with a high and low magnitude effect [high, low]: Community connectance [0.09, 0.06], Food-web trophic height [3.43, 2.05], Mass ratio Predator-Prey [1000, 10], Predator diet-breadth [12, 1] - all other model parameters are held at zero (i.e. the average effect). Estimates of community change are drawn from model predictions and describe the mean effect and 95% confidence intervals - these intervals are not visible as uncertainty is low. The stability axis is presented on a log_10_ scale. The dashed line describes the average conditions in the intact community.

Top predators frequently function as keystone species whose top-down pressure is key to regulating herbivore and mesopredator populations, which left unregulated, exert excessive pressure on lower-trophic levels, eroding ecosystem functions and triggering reduction in resilience^12,13^. This is evident in our results, with declines in richness (i.e. additional extinctions) and stability, countered with elevated biomass at lower trophic levels (Figure 1). These impacts are likely widespread in real-systems with top predators now extirpated from across much of their historical range in both marine^14^ and terrestrial^15^ systems. The decline in top predators is linked to rapid socioeconomic growth^16^, exploitation^14^, farming system and human population density^15^ - threats that continue to persist today. Preventing any additional trophic downgrading should therefore be a central objective for area- and species-based conservation.

## Predators can establish in depleted communities

Upon reintroduction, predators generally had a high probability of establishing across many scenarios (Figure 2). However, establishment success depended strongly on community intactness and on predator diet breadth. The probability of establishment (Figure 2A) dropped below 0.9 when roughly half of the community had gone extinct (community intactness: depleted) or when the predator was a dietary specialist (eating only one species). Even when reintroduction was successful, predator biomass was almost always lower than in intact communities (Figure 2B). For example, when half of the predator’s prey species were extinct, the reintroduced predator’s biomass declined by more than 20%. By contrast, when all prey species were still present, predator biomass returned to intact levels. Across all scenarios, reintroduced predator populations were generally more stable (i.e. less variable through time; Figure 2C), and greater stability is associated with a reduced risk of extinction from stochastic events^17^.

**Figure 2.**
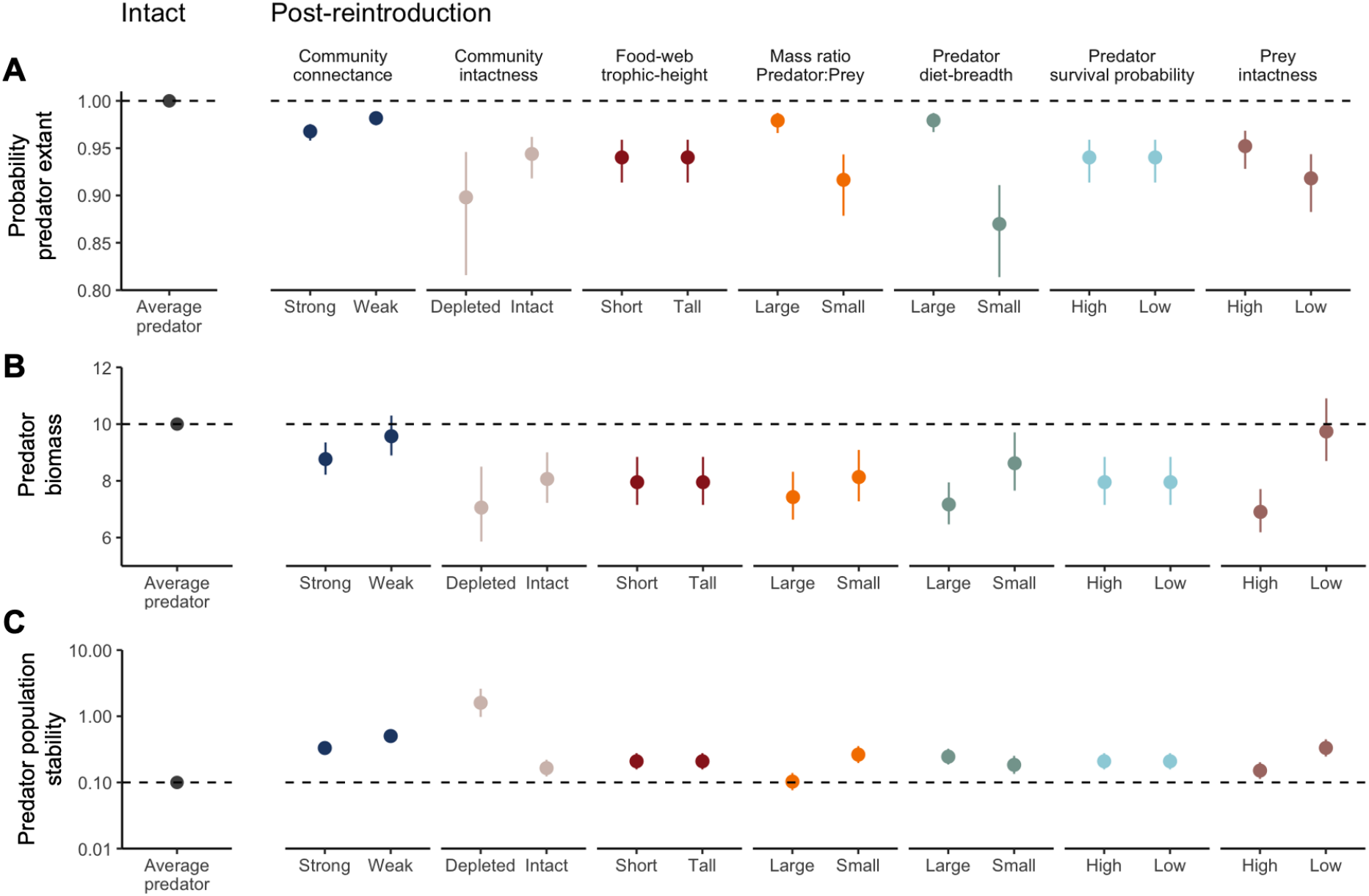
Predator responses upon reintroduction to a food-web. Change in predator status (whether the predator survives reintroduction), abundance and stability, biomass and stability (from top to bottom) in a hypothetical ‘average’ community under seven post-reintroduction scenarios each with a high and low magnitude effect [high, low]: Community connectance [0.09, 0.06], Community intactness [1, 0.5], Food-web trophic height [3.43, 2.05], Mass ratio Predator-Prey [1000, 10], Predator diet-breadth [12, 1], Predator survival probability [1, 0], Prey intactness [1, 0.5] - all other model parameters are held at zero (i.e. the average effect). Estimates of population change are drawn from model predictions and describe the mean effect and 95% confidence intervals. The stability axis is presented on a log_10_ scale. The dashed line describes the average conditions in the intact population.

Our simulations also show that predators can re-establish even in markedly depleted communities, but typically at reduced biomass relative to intact states. This seemingly counterintuitive result is consistent with empirical recoveries of large predators recolonising degraded or human-dominated landscapes via dispersal, once persecution reduces ^18,19^. Two features dominated re-establishment probability in our model: community intactness and diet breadth. Establishment remained likely when a sufficient fraction of prey species persisted and the predator was a generalist able to switch among resources; by contrast, the odds fell sharply in highly degraded webs and for dietary specialists that could not compensate for missing prey. These results suggest that specialist species in highly degraded landscapes are at greatest risk of extinction.

## Reintroduction depletes food-webs

Past evidence suggests that predator reintroduction can restore community dynamics in depleted systems^2,7^. However, our in-silico experiments reveal a more complex picture. In many cases, predator reintroduction further perturbed and degraded communities rather than restoring them. Across most scenarios, reintroductions triggered additional extinctions, regardless of the food-web structure or degree of depletion (Figure 3A). Community biomass also declined to levels lower than those observed in either intact or extirpated communities (Figure 3B), and community stability decreased after reintroduction, in some cases by as much as two orders of magnitude. One of the few exceptions occurred in communities where half of the species from the intact community were already extinct (i.e. low community intactness). In these cases, biomass and stability did not change following predator reintroduction, likely because the initial extirpation had already altered the system so extensively that reintroduction had little additional effect. Community complexity also influenced outcomes: in weakly connected food webs, predator reintroduction had little effect on biomass or stability, whereas in strongly connected food webs, reintroduction effects were more pronounced. Only one community metric, evenness (i.e. the distribution of biomass among species) improved after reintroduction, consistently to returning towards intact-like levels. The only exception was in highly depleted communities, where evenness increases dramatically, well above intact levels.

**Figure 3.**
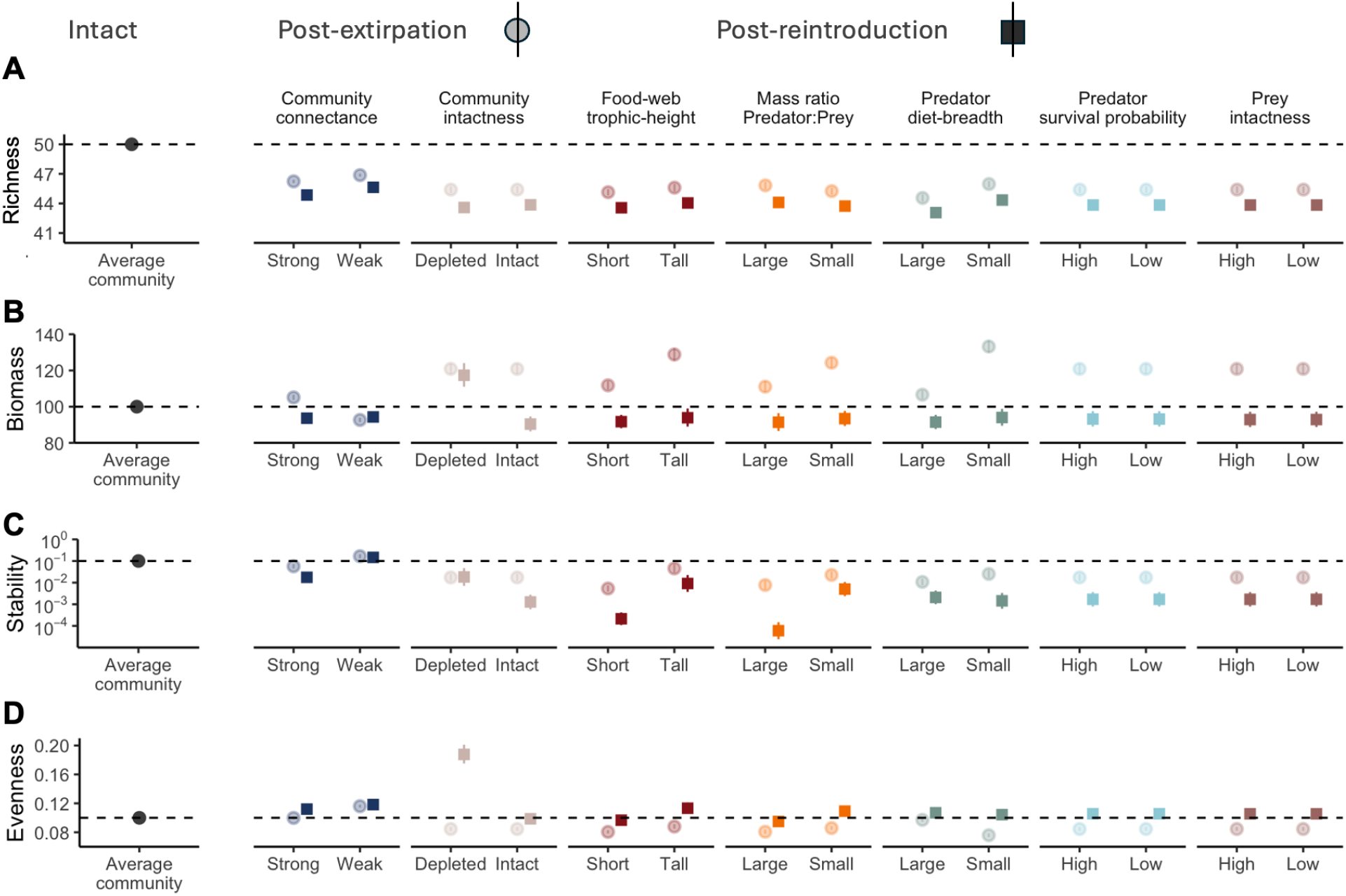
Community responses to the reintroduction of the top predator. Change in community richness, biomass, stability and evenness (from top to bottom) in a hypothetical ‘average’ community under seven scenarios each with a high and low magnitude effect [high, low]: Community connectance [0.09, 0.06], Community intactness [1, 0.5], Food-web trophic height [3.43, 2.05], Mass ratio Predator-Prey [1000, 10], Predator diet-breadth [12, 1], Predator survival probability [1, 0], Prey intactness [1, 0.5] - all other model parameters are held at zero (i.e. the average effect). We display relative community changes between the post-extirpation (transparent circle) and post-reintroduction (opaque square) transitions to emphasise any additional community change. Estimates of community change are drawn from model predictions and describe the mean effect and 95% confidence intervals - these intervals are not visible as uncertainty is low. The stability axis is presented on a log_10_ scale. The dashed line describes the average conditions in the intact community.

## Designing successful reintroductions

Conservation interventions such as species reintroduction are widely promoted as routes to bending the curve of biodiversity loss^1^, with the promise of lowering extinction risk for the reintroduced species and, in some cases, restoring wider community dynamics and ecosystem functions^2,7^. Our results suggest that, if judged solely by the fate of the focal species (as is common), many reintroductions would be deemed successful: establishment probabilities were typically high and population stability often increased, which is associated with a decreased local extinction risk^17^. Yet this species-centric view can be misleading. Across a broad set of scenarios, reintroductions triggered additional extinctions, reduced total biomass and lowered stability, indicating that population recovery can mask community-level declines.

A more reliable evaluation of reintroduction outcomes requires multidimensional success criteria that encompass composition, function and dynamics. In our simulations, community evenness increased after predator reintroduction, a positive signal that biomass was shared more evenly among species and that top-down regulation helped nudge structure toward intact-like distributions - as predicted^2,7^. However, these gains occurred alongside declines in richness, total biomass and temporal stability. Evenness should thus be treated as a useful but partial indicator of recovery. We recommend that assessments consider composition (richness, evenness), function (productivity/biomass) and dynamics (stability), so that rises in evenness are interpreted alongside metrics for functional health and stability.

Many of the most celebrated reintroduction stories - wolves in Yellowstone^2^ and sea otters in Alaska^7^ - involved keystone species whose return had far-reaching consequences for ecosystems, with impacts extending well beyond the simple predator–prey interactions we captured - a limitation of our study. However, such feedbacks, including habitat modification, nutrient cycling, or behavioural cascades like the ‘ecology of fear’, are unique features of keystones, and judging reintroduction success primarily on these iconic cases risks giving a misleadingly optimistic view. In reality, many reintroductions involve species without such dominant roles, and in these cases our results suggest that reintroductions can introduce risks to community stability and function.

To design effective reintroductions, we must ensure the community benefits from the arrival of a returning species. Theories exploring community assembly, which consider how order of arrival, dispersal, and landscape context govern which assemblages form and persist, must be integrated into reintroduction viability assessments, beyond the simple “is a food resource available?”. This is particularly true for very ambitious initiatives aiming to use reintroduction to restore heavily degraded landscapes e.g. rewilding through Lynx reintroduction. The ambition of these initiatives offering benefits to biodiversity and society, should be applauded, but will only be realised if they are designed effectively. And our results argue that reintroduction is rarely cost-free. Among all predictors, community intactness had the strongest effect on outcomes, but in a counter-intuitive way. We found that reintroductions into moderately degraded webs—those that had lost around half of their species—produced the largest gains in biomass and evenness and the smallest penalties to stability. This suggests reintroduction is perhaps most useful in degraded landscapes, but should still be applied cautiously.

Another opportunity to maximise reintroduction success is to act rapidly after extirpation. Although our simulations reintroduced predators only after the community had relaxed to a new equilibrium, earlier intervention during the transient phase could intercept trajectories before compensatory shifts are instilled - exploiting surplus prey and reducing secondary extinctions - provided the original threat has been mitigated. The importance of timing has been shown in past work^20^, but actually implementing a reintroduction at pace is challenging as we often do not know if or when a species has gone locally extinct. This is, however, becoming more feasible with modern monitoring (e.g., remote sensing, acoustic arrays, eDNA) coupled with predictive frameworks. Dynamical models, such as those used here, offer a safe test-bed to predict when a species is going to go extinct, as well as explore counterfactuals and identify scenarios where community costs are likely. Using theoretical models offers the ability to stress-test alternative strategies before implementation, although some work has argued their findings lack the realism and precision to guide actual conservation efforts^21^.

Our results also relate directly to rewilding, natural dispersal, and biological invasion. In rewilding programmes, whether releases are planned or recovery occurs via natural recolonisation, our results suggest the arrival of a strong interactor can restructure food-webs, redistribute biomass, and alter stability. This may be considered the entire objective of rewilding initiatives^3^, but without sufficient lower-trophic resources and appropriate sequencing, recolonising predators may replicate the same community-level costs we observe for managed releases. Invasive species can operate through the same mechanisms, with added risks from novelty, amplifying extinction cascades and profoundly changing food-webs^22^.

In real-world scenarios, a key obstacle to assessing whether reintroductions are truly successful is the absence of intact baselines. In simulations we know what an intact community looks like, but in reality most ecosystems are already so degraded that we lack a clear picture of intact communities. As a result, success is usually measured against a degraded reference point rather than a true intact state, creating the risk of shifting-baseline syndrome^23^. Given this challenge, we argue it is more useful to focus on transitions toward improved states than to attempt recreating past ones. There are many possible visions of what an improved state could look like - for example, the alternative pathways articulated in the Nature Futures Framework^24^ - and not all are optimised for biodiversity recovery. Regardless of which vision is pursued, two dimensions should be prioritised: (1) productivity—the capacity to generate and retain biomass, and (2) stability—the resistance and resilience of systems to disturbance. The ultimate goal is not to recreate the past communities, but transition existing communities towards an improved state.

Predator reintroduction is not a shortcut to ecosystem recovery. While focal predators often establish, their return can erode community biomass, richness and stability, masking ecological decline behind single-species success. These outcomes depend on prey persistence and food-web characteristics, factors that can be evaluated in advance with dynamic models, informed by monitoring, and integrated into planning. By reframing reintroduction as a community-level intervention rather than a demographic rescue, we provide a pathway to reduce risk in ambitious restoration programmes under rapid biodiversity change. Success should be judged holistically, with emphasis on recovering community structure, function and resilience, not solely the fate of a target species. More broadly, our finding that reintroducing predators can further destabilise community dynamics, raises a critical question: if reintroductions cannot fully restore degraded communities, are historical community states lost forever? At global scales, could this represent a fundamental constraint on attempts to bend the curve of biodiversity loss?

## Methods

We explored how population and community dynamics changed in response to predator extinction and reintroduction, examining three transition: :1) Community responses to extirpation - how intact communities change when a predator is extirpated; 2) Predator reintroduction - how predator dynamics change (relative to their intact state), when predators are reintroduced to the now degraded (because of the extirpation) community; and 3) Community responses to reintroduction - assessing how a degraded community responds to a predator’s arrival.

### Food-web simulations

We simulated food web biomass dynamics using the bioenergetic model^10,11^. The model can be defined in a system of two ordinary differential equations, one for the primary producers and one for the consumers. Primary producers gain biomass following a logistic growth and lose biomass by being fed by consumers. Consumers gain biomass by feeding on their resources (either primary producers or other consumers) and lose biomass through metabolic losses and through being consumed. The consumption rate of a consumer on a given resource increases according to the metabolic rate, the maximum consumption rate of the given consumer, and on the functional response which describes how the consumption rate of the consumer scales with its preference for the resource and its availability. The scale of the functional response is modulated by the half-saturation constant and the intraspecific consumer interference. We specified a Type 3 functional response which is controlled by the hill exponent (h = 2).

We generated food-webs using the niche model, a structural model known for creating food-web topology with similar properties to empirical food-webs^25^. To explore a range of network structures, we varied initial species richness (2–59) and connectance (0.03–0.39). Species body masses were equal to the predator–prey mass ratio (10–1000) raised to the power of the species trophic level minus one^26,27^. Metabolic rates were then determined using the allometric scaling relationship between body mass and metabolism derived from the Metabolic Theory of Ecology^10,28^, with parameter values taken from previous studies^26,27^.

The dynamics of each food-web was run in three chronologically ordered sequences. First, the food-web dynamics were run until steady state, which defined the intact state of the community. Second we extirpated the top predator of the intact community by setting its biomass to 0 (the initial biomass of the other species being set at their steady state values) and simulated the dynamics until steady state, which defined the extirpated state of the community. Third, we reintroduced the extirpated top predator in the extirpated community by setting its biomass to the average of the other species biomass, and simulated the dynamics until steady state, which defined the reintroduced state of the community. It is important to note that the predator diet preferences were not reset before each phase in the sequence, so the predator still has the capacity to consume all originally specified prey, but some prey may now be extinct, so these feeding links become inactive. We considered that biomass reached steady state when the sum of the derivative reached a 10^-6 upper threshold, and a species was considered as extinct when its biomass fell below 0. For the intact state scenario, the species initial biomass were set according to a random uniform distribution between 0 and 1. In the extirpated and reintroduced states, we set the species biomass (except for the top predator) to their steady state biomass of the previous state. The simulations were run with the default adaptative solver of the DiffEq.jl Julia package.

We collected two types of metrics from the simulations, related to topological food-web structures and to biodiversity dynamics - all defined in Table S1. For food-web structure we recorded richness (number of species present), connectance (number of trophic links divided by the number of possible links), trophic height (average trophic level of predator’s prey), diet breadth (number of resources for the top predator), mass ratio (predator mass divided by prey mass) - all measured in the intact phase. Outside of the intact phase, we also recorded prey intactness (proportion of predator’s prey species alive at the point of predator reintroduction). Our biodiversity dynamics are estimated at the population (i.e. the predator) and community scale. For the population we recorded whether the predator survived the reintroduction, as well as the the change in biomass (i.e. abundance) and population stability (inverse of the temporal variance in the predator’s biomass) between the intact and reintroduced phases. For the community we recorded the community richness, biomass, stability (i.e. inverse of the variance of the total biomass) and evenness (i.e. variability in biomass across species). These were recorded in all three phases and decomposed into relative changes between the two transitions i.e. extirpated richness divided by intact richness; reintroduced richness divided by extirpated richness. To ensure all metrics characterising biomass dynamics were independent from when the simulation was stopped (after reaching steady state), all the simulations were run a second time for 200 timesteps with initial biomass set to their steady state values. We then used the last 100 timesteps to compute biomass metrics.

### Statistical modelling

We built statistical models to characterise population and community responses to the extirpation and reintroduction. At the level of the predator, we used three models to assess the change in predator biomass, stability and probability of population re-establishment between the intact and reintroduction states. At the level of the community, we used four models to explore the change in community richness, biomass, stability and evenness at each transition (eight models total).

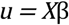

Our 11 models had the same core linear structure where the predicted response *u* is the product of a matrix of predictors *X* and associated coefficients β.

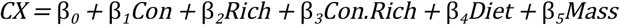

We included predictors that we recognised as important in the literature, as well as those that we considered likely relevant (Table S1). The specific predictor variables vary between the models (response variables) but all models share the same core structure *CX*, with an intercept β _0_, and five slope coefficients β: Connectance (*Con*), Richness (*Rich*), an interaction between connectance and richness (*Con. Rich*), predator diet breadth (*Diet*), and predator-prey mass ratio (*Mass*).

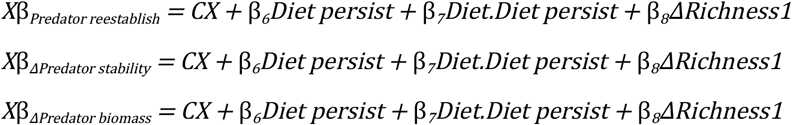

Three models are focussed on characterising the predators ability to survive the reintroduction ( *Predator reestablish*), and the change in the predators population stability and biomass post-reintroduction relative to the intact communities (*Predator stability* and *Predator biomass*). All three of these models contain two additional predictors, relative to the core predictors *CX*, describing the persistence of the community post-predator extirpation (*ΔRichness*) and the persistence of the predators prey base (*Diet persist*).

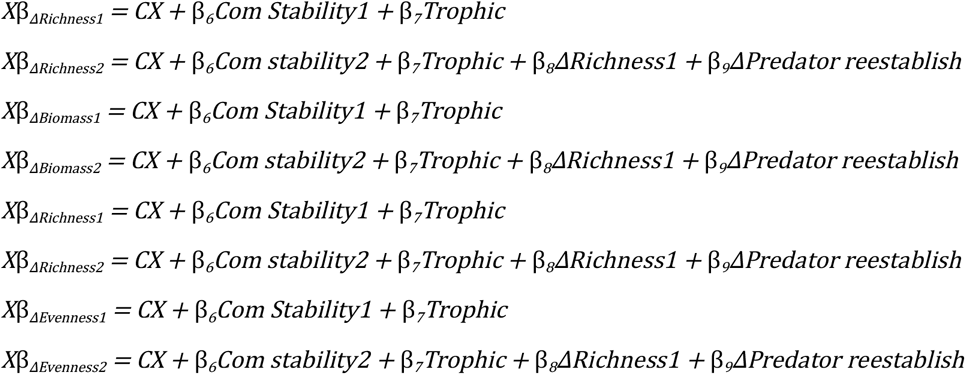

Eight models characterise the communities’ response to the extirpation (change in community between the intact and extirpated phase; numbered 1) and reintroduction (change in community between the extirpated and reintroduced phase; number 2). Specifically, *ΔRichness* describes the proportional change in richness between the phases. Biomass, stability and evenness are the relative change in community total biomass, community stability and evenness between the phases (e.g. biomass in the extirpated community divided by biomass in the intact community).

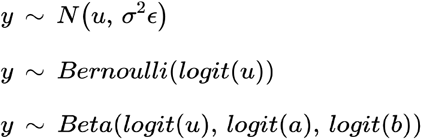

Our linear predictions from *X*β are linked to actual data through a series of general and generalised functions. All biomass, stability and evenness models are drawn from a general normal distribution with residual error σ^2^ϵ. The probability of the predator re-establishing within the community is drawn from a generalised Bernoulli distribution with a logit transformation of the linear predictor *X*β.

Persistence is drawn from a generalised beta distribution with a logit transformation of the linear predictor *X*β, and is influenced by two shape parameters αand *b*. In some cases we used a log transformation to recognise non-linearity in predictors. We also log transformed some of the response variables within the normal distribution models to correct for non-normality in residuals - all detailed in Table S1.

## Acknowledgments

TFJ was supported by a Leverhulme Early Career Fellowship ECF-2023-025. APB, TFJ and AD were supported by UKRI-NERC Grant NE/T003502/1. APB also acknowledges UKRI-NERC Grant NE/S001395/1. TH was supported by the University of Sheffield.

## Contributions

Conceptualisation: TFJ, TH, AD, APB; Methodology: TFJ, TH, AD, APB; Software: TFJ, AD; Formal analysis: TFJ, TH, AD; Data curation: TH, AD; Writing - original draft: TFJ, TH, AD; Writing - review & editing: TFJ, TH, AD, APB; Visualisation: TFJ; Supervision: TFJ, APB; Administration: APB; Funding acquisition: TFJ, APB

## Supplementary material

**Table S1.** Description of response and predictor variables used in models to characterise population and community responses to reintroduction.

| Variable | Description |
| --- | --- |
| Connectance | Measure of the food-web complexity, defined as the ratio between the number of trophic links and the number of possible links. |
| Richness | Species richness in the intact phase |
| Diet breadth | Number of resources of the target top predator (i.e. the species extirpated and reintroduced) in the intact phase |
| Mass-ratio | Predator Prey Mass Ratio: define the body mass ratio between trophic level. A value of 100 means the species of trophic level 2 have a body mass 100 times higher than the species at the trophic level 1. |
| Diet persistence | Proportion of resources of the reintroduced predator that survived in the transition from the extirpated phase relative to the intact phase. |
| Trophic level | Describe the vertical position of a species in the food-web. Computed recursively from the bottom of the food-web such as the trophic level of a species is equal to average trophic level of its prey added to one. |
| Community stability | Temporal stability of community biomass measured as the inverse of the coefficient of variation. Variable log transformed prior to modelling. |
| $P$ (Predator reestablishing) | Binary indicator of whether the predator survived its reintroduction (1: Survived; 0: Died). |
| $\Delta$ Predator stability | The change in predator population stability derived by dividing stability in the re-introduction phase by the intact phase. Variable log transformed prior to modelling |
| $\Delta$ Predator biomass | The change in biomass, derived by dividing biomass in the re-introduction by the intact phase. Variable log transformed prior to modelling |
| $\Delta$ Community stability (extirpation/intact) | The change in community stability derived by dividing stability in the extirpation by the intact phase. Variable log transformed prior to modelling |
| $\Delta$ Community stability (reintroduction/extirpation) | As above, but comparing the reintroduction and the extirpation phase. |
| $\Delta$ Biomass (extirpation/intact) | The change in total community biomass, derived by dividing biomass in the extirpation by the intact phase. Variable log transformed prior to modelling |
| $\Delta$ Biomass (reintroduction/extirpation) | As above, but comparing the reintroduction and the extirpation phase. |
| $\Delta$ Richness (extirpation/intact) | The change in species richness, derived by dividing richness in the extirpation by the intact phase. Commonly termed persistence. The predator is excluded from the richness calculation. |
| $\Delta$ Richness<br>(reintroduction/extirpation) | As above, but comparing the reintroduction and the extirpation phase. |
| $\Delta$ Evenness<br>(extirpation/intact) | The change in community evenness, derived by dividing evenness in the extirpation by the intact phase. Variable log transformed prior to modelling |
| $\Delta$ Evenness<br>(reintroduction/extirpation) | As above, but comparing the reintroduction and the extirpation phase. |

